# LITE microscopy: a technique for high numerical aperture, low photobleaching fluorescence imaging

**DOI:** 10.1101/181644

**Authors:** TC Fadero, TM Gerbich, K Rana, A Suzuki, M DiSalvo, KN Schaefer, JK Heppert, TC Boothby, B Goldstein, M Peifer, NL Allbritton, AS Gladfelter, AS Maddox, PS Maddox

**Affiliations:** Department of Biology, The University of North Carolina at Chapel Hill; Department of Chemistry, The University of North Carolina at Chapel Hill; Joint Department of Biomedical Engineering, The University of North Carolina at Chapel Hill and North Carolina State University

## Abstract

Fluorescence microscopy is a powerful approach for studying sub-cellular dynamics at high spatiotemporal resolution; however, conventional fluorescence microscopy techniques are light-intensive and introduce unnecessary photodamage. Light sheet fluorescence microscopy (LSFM) mitigates these problems by selectively illuminating the focal plane of the detection objective using orthogonal excitation. Orthogonal excitation requires geometries that physically limit the detection objective numerical aperture (NA), thereby limiting both light-gathering efficiency (brightness) and native spatial resolution. We present a novel LSFM method: **L**ateral **I**nterference **T**ilted **E**xcitation (**LITE**), in which a tilted light sheet illuminates the detection objective focal plane without a sterically-limiting illumination scheme. LITE is thus compatible with any detection objective, including oil immersion, without an upper NA limit. LITE combines the low photodamage of LSFM with high resolution, high brightness, coverslip-based objectives. We demonstrate the utility of LITE for imaging animal, fungal, and plant model organisms over many hours at high spatiotemporal resolution.

## Introduction

To properly visualize and measure cellular and subcellular dynamics, cell biologists demand imaging at high spatial and temporal resolution. The fluorescence microscope is a popular modern tool used to address these demands and solve cellular dynamics problems. However, conventional fluorescence microscope modalities require high intensity light to illuminate the sample through the objective lens, exciting all fluorophores in the path of the collimated excitation light. The fluorophores emit light that is collected by the objective lens and transmitted to the detector. A disadvantage of the traditional “epi-illumination” geometry is that light is emitted from fluorophores outside the focal plane and contributes to the image, which confounds the focal information. Confocal microscopy mitigated this problem by selectively collecting light from the focal plane through the use of conjugate pinholes^1^. However, the reduction of out-of-focus fluorescence by confocal microscopy does not overcome the need for high-intensity illumination light that generates out-of-focus excitation events (Fig. 1A; blue box). High intensity illumination transmits intense energy to the sample, damaging fluorophores that release reactive oxygen species upon photobleaching. Consequently, these reactive oxygen species chemically damage living samples through phototoxicity^2^.

**Figure 1.**
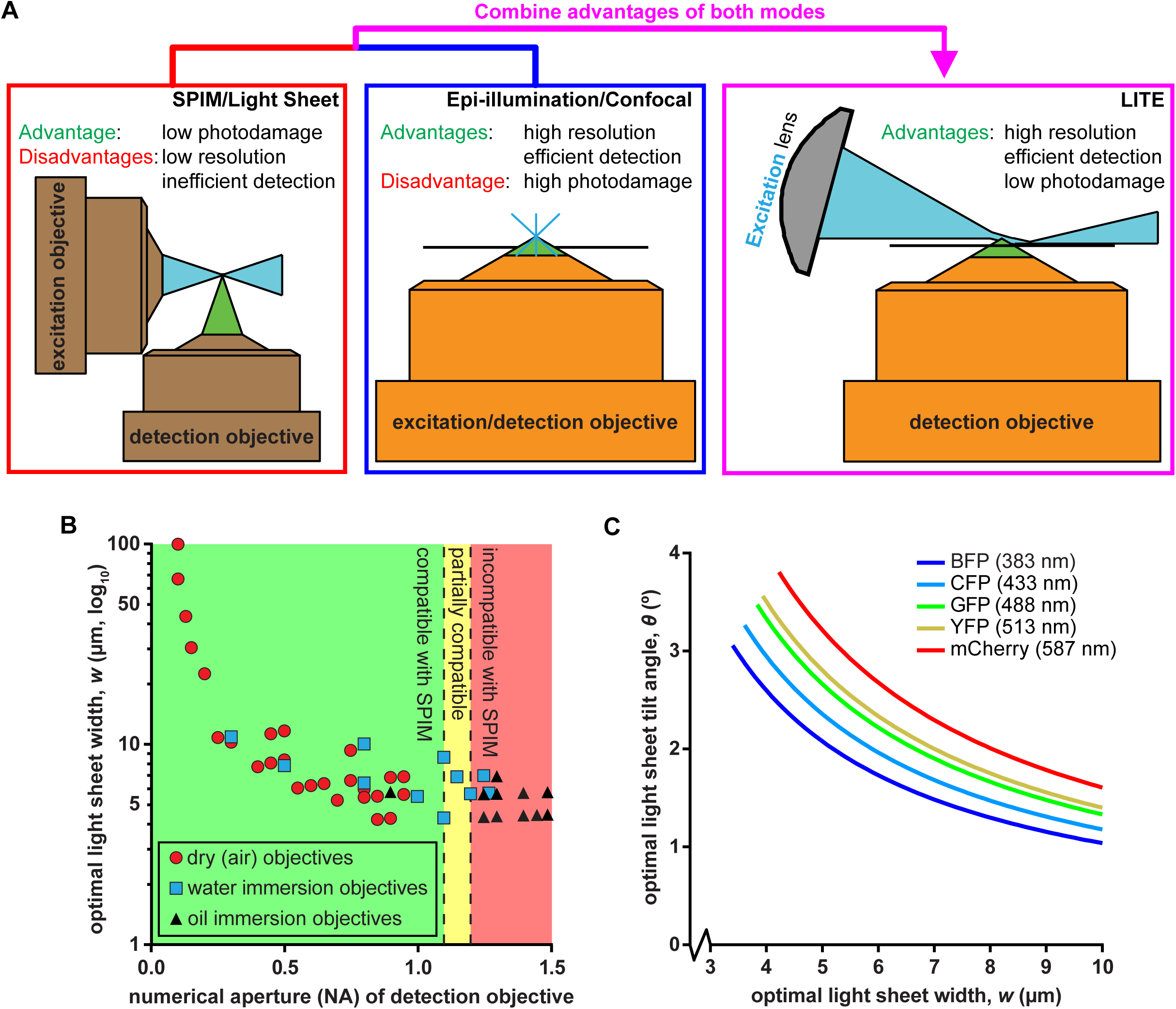
Rationale and theory behind LITE. (1A) We combined low photodamage of SPIM/LSFM (left) with high-NA objectives (orange) of Epi-illumination/Confocal microscopy (center) to create LITE (right). LITE tilts a cylindrical lens (gray) to focus a laser (blue) into a sheet onto coverslip surface (black line). (1B) Scatterplot of calculated optimal light sheet width for LITE, based on equation (7), for 90 commercially available objectives (plotted by increasing NA). Green region: objectives that can/have been used with existing SPIM technologies. Yellow region: objectives that have been used with LSFM by means of unconventional geometries^5^. Red region: objectives previously incompatible with SPIM/LSFM. (1C) Theoretical optimal light sheet width, ***w***, as a function of optimal sheet tilt angle, ***θ***. Ideal light sheet parameters are traced for five common fluorescent proteins: blue fluorescent protein (BFP), cyan fluorescent protein (CFP), green fluorescent protein (GFP), yellow fluorescent protein (YFP), and monomeric Cherry (mCherry). Wavelengths of excitation light plotted in (1C) correspond to maximum absorption wavelength of each protein.

**L**ight **S**heet **F**luorescence **M**icroscopy (LSFM, or **S**elective **P**lane **I**llumination **M**icroscopy SPIM) minimizes excitation-based photodamage by only partially illuminating the sample^3^. In the 15-year existence of modern LSFM, various implementations have arisen, most of which use two traditional objective lens elements arranged orthogonally in order to 1) illuminate the sample with a sheet of light and 2) align the detection focal plane with the illuminating sheet^3-6^. LSFM reduces or eliminates out-of-focus excitation, increasing the signal-to-background ratio (SBR) for fluorophores in the focal plane (Fig. 1A; red box). This higher SBR allows detection of image features with lower excitation energy, thus reducing the photodamage incurred with conventional optical configurations. These features allow the acquisition of a significantly larger number of exposures of a sample than any other mode of fluorescence microscopy. However, the orthogonal orientation of the illumination light sheet with respect to the detection objective generally requires that the sample be mounted at a minimum of one millimeter from the detection objective, forcing use of low numerical aperture (NA, below 1.1) detection objective lenses. Therefore, the use of highly efficient, high-resolution oil-immersion objectives is incompatible with current LSFM regimes.

The detection of subcellular structures that drive cell biological processes including mitosis, endocytosis, and cytokinesis require high-NA detection objectives, due to their increased resolution and detection efficiency. Because 1.1 was the highest feasible NA detection objective used with traditional geometries^5^ (Fig. 1B; green box), use of LSFM to study these sub-cellular structures with the traditional resolution or efficiency was not possible. Multi-view SPIM geometries have been able to accommodate a 1.2 NA water-immersion objective to increase the resolution and detection efficiency of LSFM^6^ (Fig. 1B; yellow box); however, in order to approach the native resolution of oil-immersion objectives (Fig. 1B; red box) traditionally used in cell biology, post-acquisition deconvolution was required. This data processing has high requirements for time, user expertise, specialized software, and data storage, which are currently inaccessible to the average cell biology laboratory. Accordingly, there existed a need to build upon the currently available designs for LSFM by combining selective illumination with conventional microscope stands and objective lenses that enable detection and resolution of subcellular structures and dynamics.

Here we present **L**ateral **I**nterference **T**ilted **E**xcitation (LITE) microscopy, which we developed in order to use high-NA, oil-immersion objective lenses to image samples illuminated by a light sheet (Fig. 1A, magenta box). We achieved this goal by using a tilted sheet that can access the working distance of high-NA oil- and water-immersion objective lenses, including a 60X 1.49 NA oil-immersion objective that accepts 88% more emitted fluorescence and offers a 26% increase in native lateral resolution compared to a 25X 1.1 NA water-dipping objective^5^. LITE is compatible with traditional coverslip-based mounting conditions, meaning that LITE can be used with water- and oil-immersion objectives. The LITE method can also be implemented unobtrusively on most existing upright or inverted microscope systems, meaning high-resolution differential interference contrast (DIC) or other microscopic modalities can be used simultaneously (or in rapid succession) with LITE imaging. LITE images do not require computational reconstruction to view; the native images received from the camera are the data. In sum, LITE microscopy combines the low photodamage of LSFM with the high-NA objective lenses to allow high spatiotemporal imaging.

## Methods

LITE is a novel method for introducing a light sheet within the working distance of high-NA objective lenses (Fig. 1A). Briefly, these goals were accomplished by first directing a collimated beam of excitation light through a photomask and cylindrical lens. The cylindrical lens focused the excitation light to form a roughly “wedge-shaped” beam of light. The beam converged to its minimal thickness and formed the light sheet at the focal plane of the cylindrical lens, approximately three centimeters away from the cylindrical lens. The photomask was used to pattern the focusing beam so that the light sheet was lengthened^7^. To access the working distance of high-NA lenses, the excitation light was tilted such that the bottom of the converging “wedge” was parallel to the detection objective focal plane. Thus, the light sheet was formed at the focal volume of the detection objective, in which the fluorescent sample was mounted. LITE allows mounting samples on coverslips, provided the chambers also have an optically clear opening to allow access by the converging illumination light. We have engineered several suitable chambers and present imaging data from a diverse range of model organisms.

### 1. Illumination

LITE imaging requires collimated, radially symmetric, coherent illumination light. We generated such a beam using a collimator illuminated by a laser combiner (Agilent Monolithic Laser Combiner 400, MLC 400) with an FC/APC fiber-coupled laser output of four wavelengths (405, 488, 561, and 650 nm). The four laser sources were solid state and pre-aligned to deliver a radially symmetric, coherent beam (Fig. S1). The maximum power outputs, after the fiber, of the four lasers in order of increasing wavelength were 18, 52, 55, and 37 mW, although only a fraction of each beam is used to generate the light sheet. The choice of illuminator should be based on specific application, fluorescent proteins *in vivo* in this case. An internal acousto-optical tunable filter (AOTF), analog-controllable via DAQ Board interface, was used for modulating wavelength intensities. For brevity, we mainly describe our setup as monochromatic illumination at 488 nm excitation (for EGFP).

### 2. Beam Conditioning

LITE illumination involves conditioning from the laser source such that the diameter of the radially symmetric beam is magnified to a value that is equal to or greater than the full aperture of the slits of a customized photomask (see below, Methods Part 3). The beam should remain collimated after conditioning. Here, collimation and beam expansion were combined by an FC/APC-coupled (fiber connector/angled physical contact) TIRF (total internal reflection fluorescence, Nikon Instruments) microscopy collimator that achromatically collimated the lasers to a beam diameter of 22 mm (Fig. S1).

### 3. Photomask/Cylindrical Lens System

We used a cylindrical lens to focus a radially symmetric, collimated beam along one axis in order to approximate a non-diffracting “sheet” of light at the focus of the cylindrical lens. The sheet itself (in the focus of the cylindrical lens) can be approximately defined as a rectangular prism with three dimensions: the thinnest, diffraction-limited vertical width (***w***) that the converging laser reached at the cylindrical lens focal plane, the axial length (***L***) over which the laser remained at its diffraction-limited width before diverging, and the unfocused horizontal breadth (***b***) of the laser. The full width at half maximum intensity (FWHM) of the sheet (hereafter referred to as ***w***) is defined by:

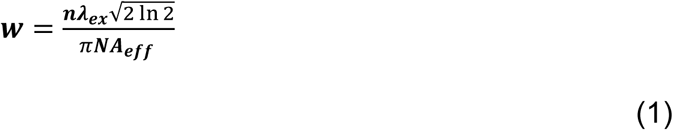

In equation (1), ***n*** is the refractive index of the medium in which the laser was focused to a sheet (typically ∼1.33 for aqueously media, although this value varies based on the temperature and chemical composition of the media, and the wavelength of the excitation light), ***λex*** is the wavelength of the excitation laser (in μm), and ***NA*_*eff*_** is the effective numerical aperture of the cylindrical lens. Note that ***NA*_*eff*_** can be smaller than the reported NA of the cylindrical lens, as ***NA*_*eff*_** depends on the percentage of the cylindrical lens NA that is used (i.e. the vertical height of the collimated excitation light incident on the cylindrical lens back aperture). Thus, ***w*** is inversely proportional to the diameter of the collimated beam incident to the cylindrical lens, assuming the beam diameter is less than the full cylindrical lens back aperture. The thinnest sheet possible is traditionally preferable in LSFM, for two reasons: (1) to minimize out-of-focus excitation/emission in the fluorescent sample and (2) to prevent photodamage in out-of-focus planes. However, the choice of sheet thickness in LITE was complicated by the mathematical interdependence of ***w*** and ***L***, in equation (2):

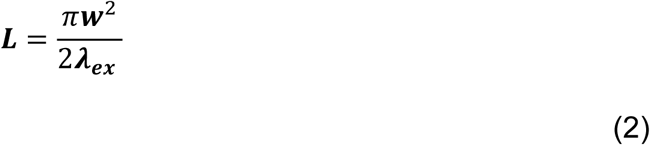

As shown in equation (2), it is evident that ***L*** increases with the square of ***w.*** Practically, this meant that the thinnest sheet possible (minimal ***w***) was not necessarily the best sheet for LITE, as the distance over which the sheet remains diffraction-limited (***L***) could have been too short to cover the field-of-view (***FOV***) of the detection objective used for detecting the signal. If the sheet began to diverge over the ***FOV***, then the illuminated slice of the fluorescent sample would vary significantly in both thickness and illumination intensity along the ***FOV***. This would result in inconsistent excitation of fluorophores, making quantitative analysis of fluorescent images difficult.

In order to maximize the ***L*** for a given ***w***, we placed a quadruple-slit photomask (FrontRange Photomask) in the principle plane of the cylindrical lens, before the beam enters the lens (Fig. S1). The theoretical and practical design of these slits were first described and implemented by Golub *et al*. in 2015. Briefly, this method increased ***L*** of a cylindrical lens-based light sheet beyond what equation (2) predicts by creating an interference pattern at the cylindrical lens focal plane between two harmonic cosine waves^7^. Golub *et al*. (2015) presented the equation for the depth of field of the elongated light sheet below in equation (3):

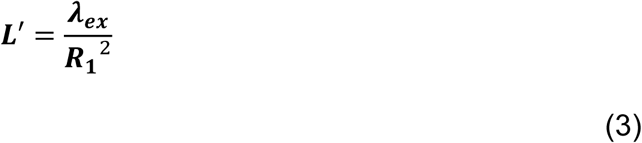

In equation (3), ***L’*** is the elongated sheet length, and ***R1*** is the radius of the inner photomask slits^7^. In order to put equation (3) in terms of ***w***, we equated ***R1*** to ***NA_eff_*** using equation (1) and substituted the equivalence into the equation (3) denominator to arrive at equation (4):

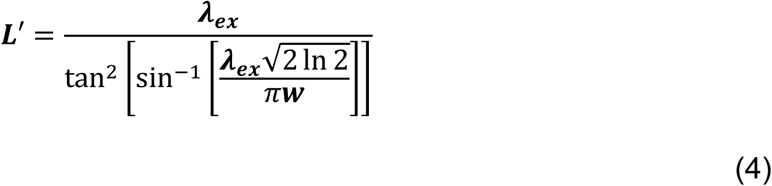

In LITE as described here, the thickness and spacing of the photomask slits were scaled from the values for a 152-mm focal length cylindrical lens^7^ to the scale of our selected 40-mm focal length, aspheric, cylindrical lens (ThorLabs; AYL5040-A). The optical trade-off of this interference strategy was the generation of side lobes and loss of illumination intensity. Side lobes should theoretically manifest as coplanar light sheets above and below the bright center peak of the main light sheet. However, more than 80% of the total laser energy should in the center sheet^7^. Side lobe minimization is important to reduce the probability of excitation and emission outside the detection objective focal plane.

### 4. Optimization of Sheet Dimensions and Parameters

Creating a non-diffracting light sheet of a width within an order of magnitude of the wavelength of light requires that the light be focused. Accordingly, previous light sheet fluorescence microscopes have used standard (or custom) objective lenses to focus a beam to create a light sheet of a minimal width in the sample^3-6^. This orthogonal, two-objective method sterically limits the choice of detection objectives to those with a long enough working distance (greater than one millimeter) to focus on the sheet, since the illumination and imaging objectives cannot touch. Here, we present a novel solution for using virtually any existing microscope objective, including those with high NA, for imaging fluorescence signal from a light sheet (Fig. 1A). This represents a significant advance in LSFM, as biologists are no longer limited in their choice of objectives (Fig. 1B). A detailed, a step-by-step method for selecting the ideal setup of a LITE microscope illuminator based upon the desired objective is presented below.

For effective imaging with LITE, it is necessary to illuminate an objective’s volume-of-view (VOV) while minimizing illumination outside the VOV. An objective’s VOV can be defined by the product of the two-dimensional field-of-view (***FOV***) and the one-dimensional depth-of-field (***DOF***). The ***DOF*** of an objective, otherwise known as axial resolution, is a set parameter that varies based on the **NA** and wavelength of the emitted fluorescence (***λ_em_***) that is collected by the objective (see equation (6) below).

The relationship between the light sheet FWHM ***w*** and the objective ***DOF*** was derived from the necessity to form the light sheet at the coverglass surface so that it is within the working distance of high-NA objectives. Confined by this geometry, it is impossible to form a light sheet that is completely orthogonal to the focal plane of a high NA objective within its standard working distance (typically < 300 μm) while also projecting the converging beam over a flat surface, such as a coverslip. To solve this problem, we tilted the collimated beam, photomask, and cylindrical lens relative to the surface of the objective. Tilting in LITE was done at a precise, but customizable, angle: the half angle of the laser as it converges in aqueous media. Tilting the LITE setup at this half angle, ***θ***, allowed the bottom part of the converging beam to propagate parallel to the coverslip surface without reflection or refraction of the laser through the coverslip before the laser reached the sample (Fig. 1A). Tilting a light sheet relative to the objective’s ***FOV*** is not typical of other light sheet modalities^3-6^. Due to this aspect of our design, it is distinct from current LSFM methods.

In order to determine the ideal light sheet dimensions (***w***, ***L***, and ***θ***) from the parameters of any desired objective (magnification ***M***, ***NA***, ***FOV***, and ***DOF***), several basic mathematical relationships were considered. We first determined a useful relationship between ***w*** and an objective’s ***FOV*** and ***DOF***. To calculate the full ***FOV*** of the desired objective, equation (5) was considered:

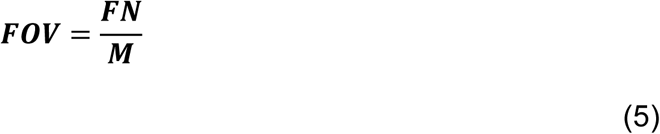

***FN*** is the field number of the objective (in μm) and ***M*** is the lateral magnification of the objective (dimensionless). Thus, ***FOV*** is the full one-dimensional diameter (in μm). If a shorter ***FOV*** were desired (i.e. the length of a camera’s pixel array), it could be instead defined manually as some fraction of the full objective ***FOV***. Next, to calculate the theoretical ***DOF*** of a desired objective, equation (6) was considered:

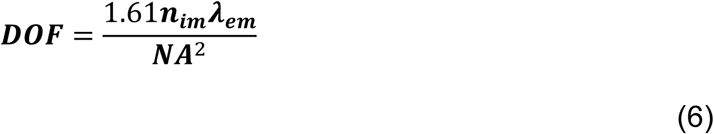

The variables in equation (6) (***n*_*im*_**, ***λ_em_***) correspond respectively to the coverslip immersion medium refractive index and fluorophore peak emission wavelength (in μm). It is worth noting that the useful ***DOF*** of an objective (axial resolution) changes based on the desired fluorophore, since both ***λ_em_*** and ***n*_*im*_** vary based on the fluorophore.

Once the ***DOF*** and ***FOV*** have been correctly identified for the objective of choice, the ideal width of the light sheet was calculated. The following equation was derived to determine the dependency of ***w*** on ***DOF***, ***FOV***, and ***λ_ex_***:

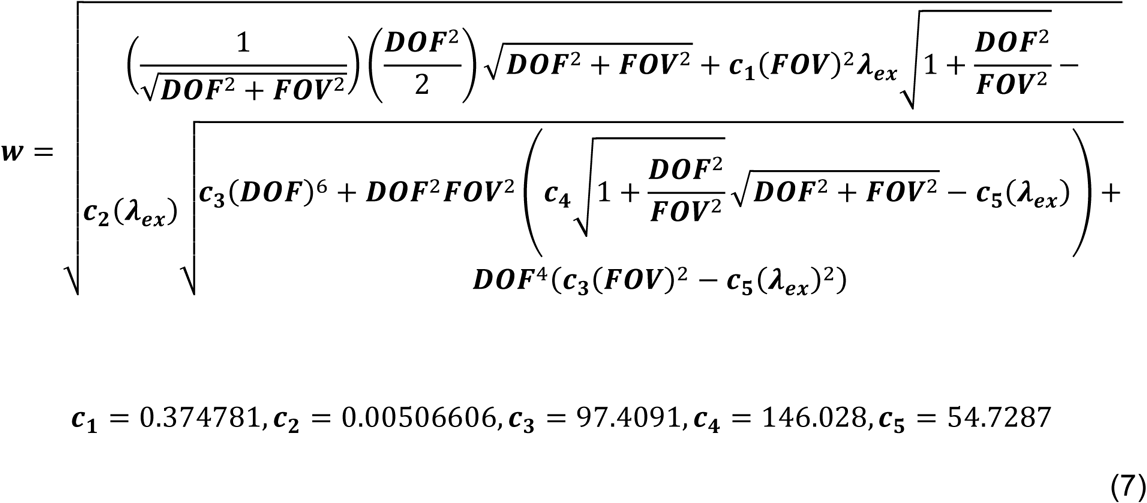

Constants in equation (7) (c_1_ – c_5_) are unchanging factors that result from the explicit derivation of ***w***. Thus, we arrived at a function of two variables such that ***w*** = *f*(***FOV, DOF***). In sum, the ideal light sheet width for any given objective could be calculated. In order to illustrate the general trend of how ***w*** varied as a function of objective parameters, we obtained the ***FOV***, ***M***, ***NA***, and ***DOF*** of 90 commercially-available detection objectives and plotted the optimal light sheet thickness (***w***) for each objective as a function of its ***NA*** (Fig. 1B).

Once the width of the light sheet was known, we then calculated the length ***L’*** over which a light sheet of that width remains non-diffracting from equation (4)^7^. Finally, we also calculated the half-angle of the converging laser, ***θ***, that forms the light sheet, through substituting the general equation for lens numerical aperture, equation (8), into equation (1) and solving for ***θ*** to yield equation (9):

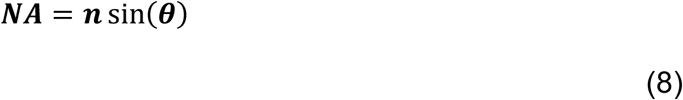

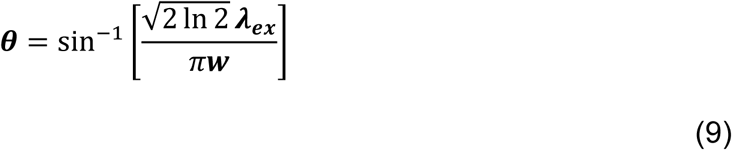

The resultant angle from (9) is the maximum angle at which the focused sheet should be tilted relative to the VOV inside the sample chamber. Our selected cylindrical illuminating lens was a dry lens that focuses the laser into air (n ∼ 1), so the laser must first refract into the sample chamber (n ≥ 1.33) before reaching the sample (see Methods Part 5 below). Equation (9) is plotted in Figure 1C in order to visually illustrate that ***θ*** decreases exponentially as ***w*** increases. Since ***θ*** and ***w*** vary with respect to the different excitation and emission wavelengths among fluorescent proteins, five traces are shown in Figure 1C that correspond to five commonly used biological fluorophores (BFP, CFP, GFP, YFP, and mCherry) and their respective maximal excitation wavelengths (383, 433, 488, 513, and 587 nm).

If the tilting is kept to the minimum ***θ*** necessary to completely illuminate the ***FOV*** of interest, then out-of-focus excitation was still dramatically reduced (compared to conventional illumination) in the case of all objectives over a wide range of numerical apertures, magnifications, and depths of field (Fig. 1B,C). A byproduct of this scheme was that ***w*** was always wider than the DOF, a feature that (in principle) lead to increased outof-focus excitation compared to conventional light sheet illumination. However, in part due to the optical sectioning ability of high-NA lenses and the Gaussian nature of light sheet intensity, this effect was not observed in practice (see discussion).

### 5. Sample Chambers

To be compatible with LITE, sample chambers must meet two main criteria: (1) have a glass coverslip as the bottom surface for use with high-NA objectives, and (2) have a flat, optically clear, and homogenous side in order to allow the laser to focus inside of the chamber at the coverslip surface. Images presented in this paper were acquired using one of two types of chambers that meet these criteria.

The first type of chamber (Fig. S2-A, hereafter referred to as Chamber A) consisted of an open-topped, media-filled box formed using four 22 x 22 mm #1.5 crown glass coverslips and one 24 x 60 mm #1.5 crown glass coverslip. The coverslips were cemented in place using VALAP (1:1:1 mixture w/w of vaseline, lanolin, and paraffin). In order for the laser to enter the chamber normal to the front coverslip surface, it was necessary to angle the front coverslip at the sheet convergence angle, ***θ***. Advantages of Chamber A included short fabrication time (∼3 minutes) and low cost per unit. Disadvantages of Chamber A included incompatibility with samples less dense than their media (samples do not sit on the surface of the coverslip while immersed in media), incompatibility with upright microscopes, irreproducibility of the tilt angle of the front coverslip, irreproducibility of sample positioning, and potential VALAP leaking into the chamber that interferes with the converging laser.

To overcome some of the disadvantages of Chamber A, a second chamber was created (Fig. S2-B, hereby referred to as Chamber B). Chamber B was a microfluidic chamber consisting of a 24 x 60 mm #1.5 glass coverslip and an imprinted piece of polydimethylsiloxane (PDMS, commercially available as Sylgard 184 from Corning; see Figure S2-C for a two-dimensional view of the imprint pattern). Two copies of the imprint pattern were copied onto a photomask for photolithography (FrontRange Photomask). Templates for microfabrication were made by spin coating 1002-F negative photoresist^8^ onto clean 50 x 75 mm glass slides at various speeds (1500-3000 rpm) for various thicknesses of photoresist (7-50 μm). Slides were exposed to 400-600 mJ of UV radiation under the patterned photomask. Unpolymerized photoresist was removed chemically, leaving behind hardened microfeatures on the template that act as a negative for imprinting PDMS. The template was placed inside a sealed metal casting chamber with two polished, flat metal sides, tilted at ***θ*** relative to the normal of the template’s surface. Pre-mixed liquid PDMS (1:10 w/w ratio of crosslinker to base) was cast over the template before vacuum de-gassing. PDMS was heated to 40 °C and left to polymerize for 24 hours before separation from the template. We used a 1.0 mm biopsy punch to create inlet and outlet channels through the PDMS into the microchamber for flowing in media/samples. PDMS molds were cleaned with methanol, ethanol, and distilled water, and coverslips were cleaned with isopropyl alcohol and acetone before both the molds and the coverslips were plasma treated for 30 seconds. PDMS molds were bonded by physical adhesion to coverslips to create the finished Chamber B. Advantages of Chamber B included high reproducibility among chambers (in shape, size, and tilt angle ***θ*** of the side), high customizability, and the ability to mount any sample in a reproducible location (close to the coverslip, in the focused sheet), compatibility with samples of low density (e.g. *Ashbya gossypii*). The main disadvantage of Chamber B was a longer production time per unit (24 hours).

### 6. Microscope hardware parameters

LITE microscopy can be used on a broad diversity of microscope stands (inverted or upright), with any objective, and with any coherent, collimated laser source. The physical setup of our LITE prototype is detailed below, but it may be easily adapted for different existing microscope hardware. The LITE apparatus was constructed adjacent to a TE2000 inverted stand (Nikon Instruments, Fig. S3). The stand is equipped with a XY motorized stage (50 mm travel, Prior Instruments) for positioning of samples and a piezo motorized Z stage (100 μm travel, Prior Instruments) for scanning the sample through the light sheet/focal plane during multi-plane acquisition.

Several custom parts necessary to position the photomask, cylindrical lens, and collimator at the appropriate angle relative to the detection objective were designed using AutoCAD for Mac 2015 (AutoDesk) and were either manufactured using a 3D printer or machined from aluminum. Computer-assisted design (CAD) files of custom parts are available from the authors upon request.

For fluorescence detection/magnification, a variety of detection objectives were used. All objectives used in this article for fluorescent organism visualization are coverslip-based, water or oil immersion, NA ≥ 1.2, infinity-corrected, with magnifications between 40 and 100X. Specific objective parameters for individual image sets are listed in the figure legends. A 535/50 nm emission filter was installed in the infinity path of the objective to filter out scattered 488 nm excitation light (Fig. S1). No other filters (e.g. dichromatic mirrors) are necessary in LITE. Magnified images were re-focused with a 1X tube lens onto an Andor Zyla 4.2 sCMOS camera. Laser AOTF (Methods Part 1), motorized Z piezo position, and camera firing were trigged through a DAQ board interface and controlled through Nikon NIS Elements.

### 7. Sample preparation

All *C. elegans* specimens were cultured on nematode growth media (NGM) + 2% agar petri dishes and fed with OD421 bacterial cultures over three days at 20 °C. Adult *C. elegans* were dissected in M9 media (17 mM K_2_HPO_4_, 42 mM Na2HPO4, 85 mM NaCl, 1 mM MgSO_4_) to obtain embryos, which were mounted in Chamber A with M9, or M9 + 2 mM NaN_3_ (for photobleaching measurements). Strains used include LP148 (*unc-119(ed3) his-72(cp10[his-72::gfp+ LoxP unc-119(+) LoxP]*) III)^11^ and LP447 ((cp178[klp-7::mNG-C1^3xFlag]) III, unpublished).

HeLa cells stably expressing Hec1-EGFP were cultured in Dulbecco’s modified Eagle’s medium (DMEM: ThermoFisher Scientific) supplemented with 10% fetal bovine serum (FBS: Sigma), 100 U mL^-1^ penicillin and 100 mg mL^-1^ streptomycin at 37 °C in a humidified atmosphere with 5% CO_2_. Rounded mitotic cells were shaken off one hour prior to imaging and mounted in L-15 medium (ThermoFisher Scientific) in a poly-L-lysine-coated (Sigma) Chamber A. Cells were kept at ∼32 °C while imaging.

*A. thaliana* seeds were surface sterilized and plated onto 0.5X Murashige and Skoog salts^9^ and 0.6% Phyto Agar (Research Products International) plates. Germination was induced by incubation at 4 °C for 48 hours, when plates were moved to a growth incubator equipped with a mix of fluorescent and incandescent lights and set to 23 °C. After five days, seedlings were mounted in Chamber A coated with poly-L-lysine-coated (Ted Pella) and covered with a thin slab of 2% agarose before the chamber was filled with distilled water.

*D. melanogaster* adults were mated for three days at 25 °C on apple juice plates. Embryos were collected 3-5 hours post egg-laying, and dechorinated in 50% bleach for five minutes. Embryos were then sorted under a dissecting scope, looking for embryos at pre- or early-germband extension. Embryos were then mounted on a poly-L-lysine (Ted Pella) coated coverslip and coated with halo-carbon oil #700 (Lab Scientific) in Chamber A (Fig. S2-A) for imaging. The following stocks were obtained from the Bloomington Stock Center: *Maternal alpha tubulin GAL4* (7062), and UAS-Axin:GFP (7225).

*Hypsibius dujardini* tardigrade cultures were maintained in 2L flasks with oxygenation using spring water (Poland Spring) as culture media and fed *Chlorococcum sp*. algae. To isolate specimens, a small amount of culture was decanted into a 60 mm Petri dish and animals were transferred to a 1.5-mL microcentrifuge tube using a dissecting microscope and mouth pipet. Specimens were fixed as previously described in Smith and Jockusch (2014)^10^. Briefly, specimens were relaxed in carbonated water for hour before fixation. Specimens were fixed in 4% EM Grade Paraformaldehyde (Electron Microscopy Sciences) in PB-Triton (1X phosphate-buffered saline, 0.1% Triton X-100, pH 7.4) for 15 minutes at room temperature. Specimens were used immediately for staining. Fixed specimens were stained with phalloidin and wheat germ agglutinin. Wheat germ agglutinin (WGA) labels the cuticle of *Hypsibius dujardini* (unpublished observation, T.C. Boothby). Alexa Fluor 594 WGA (Thermo-Fisher) was diluted to 10 μL/mL in PBS and specimens were incubated in this solution overnight. Phalloidin staining was performed as described in Smith and Jochusch (2014)^10^. After WGA labeling, specimens were washed four times for 15 minutes and then left overnight in PB-Triton with 0.1% NaN_3_. Specimens were incubated for a day in a 1:40 dilution of phalloidin (Oregon Green 488 conjugated - Molecular Probes) in PB-Triton with 0.1% NaN3 and then washed three times for five minutes in PB-Triton. Fixed and stained tardigrade adults were kept at 4 °C until imaging. Tardigrades were moved to a PBS-filled Chamber A and positioned using mouth pipette for imaging.

*A. gossypii* cells were germinated in Chamber B at 30 °C for eight hours in *A. gossypii* 2X low fluorescence media (YNB +N, -folic acid, -riboflavin (Sunrise Scientific, 1.7 mg/mL), CSM-ade (1.6 mg/mL), *myo*-Inositol (1.0 mg/mL), dextrose (20 mg/mL), aspartic acid potassium salt (7.0 mg/mL), glutamic acid potassium salt (7.0 mg/mL), adenine hemisulfate (10 μg/mL), pH 7.0) with ampicillin (100 μg/mL) and G418 (200 μg/mL). After eight hours, fresh media was added and chambers were moved to room temperature for imaging.

### 8. Image processing

Images were acquired using NIS-Elements (Nikon Instruments). Unless otherwise specified, all images presented in this paper are raw acquisition data (after camera offset subtraction). No post-acquisition deconvolution or stitching is required to view LITE images, although for some multi-plane movies have been maximum intensity projections (max IP) were generated in the z dimension or deconvolved (specified in the figure legends). Fluorescence intensity measurements, kymographs, maximum intensity projections, image scaling, false-coloring, and movie annotations were performed using Fiji. Richardson-Lucy deconvolution images and three-dimensional supplementary movies were made using NIS-Elements (Nikon Instruments).

## Results

### LITE illuminates a thin slice of fluorescent samples

The feature shared by all SPIM/LSFM technologies is the spatial restriction of the illumination light to a volume on the order of magnitude of the detection objective’s focal plane, so that fluorophores outside of the focal plane do not experience unnecessary illumination. We used LITE microscopy to produce a sheet of light with constant thickness over the desired objective’s field of view (150 μm). We theorized we could accomplish this thin illumination scheme using established cylindrical lens-based cosine wave optics^7^. We therefore calculated the theoretical side view of the light sheet to visualize the predicted sheet width and length (width = 4.3 μm, length = 300 μm; Fig. 2A). In order to verify that our experimental light sheet recapitulates what our calculations predict, we visualized the experimental sheet from the side at 1X magnification through a dilute solution of fluorescein (Fig. 2B, upper). We acquired a 40X magnified image of our experimental light sheet in order to quantify the width (Figure 2B lower, red box). When compared to the theoretical intensity profile^7^ (Fig. 2A) predicted by the theoretical electric field amplitude at the focal plane of the masked cylindrical lens, our experimentally observed central peak has nearly identical sheet dimensions (width = 4.3 μm; length = 296 μm; Fig. 2D). Practically, we observed that excitation intensity was too low in these side lobes to generate signal in low density fluorophore regimes, such as those of live cell imaging (data not shown).

**Figure 2.**
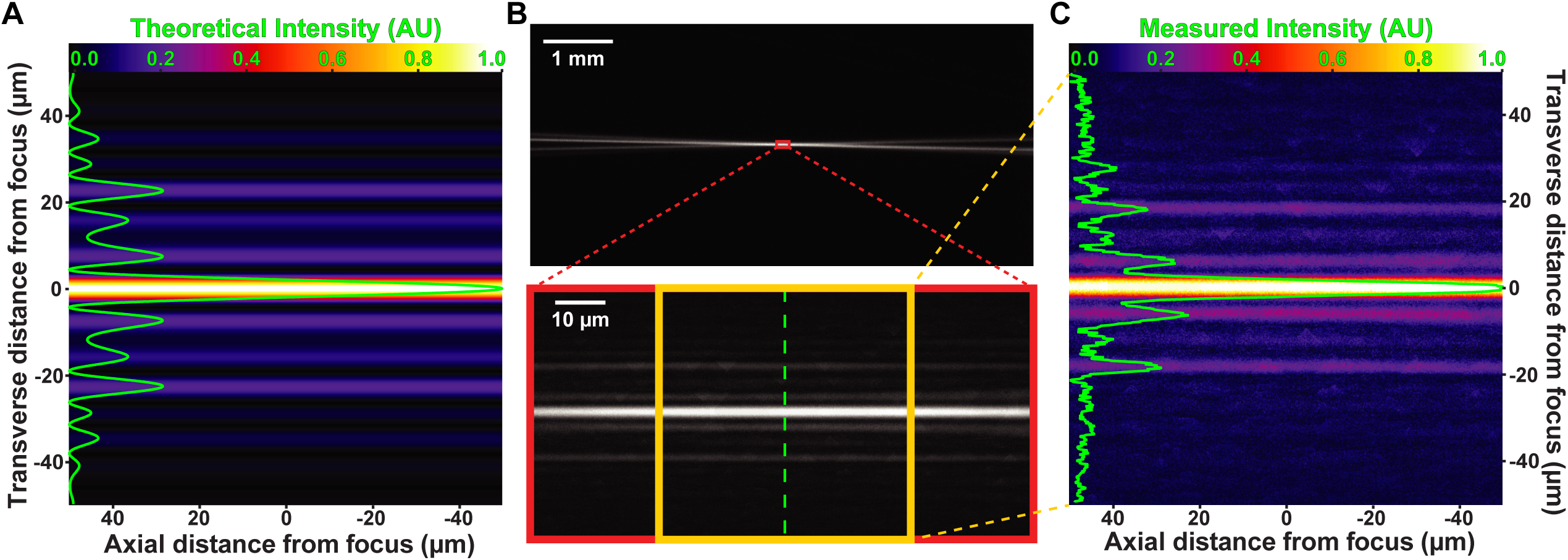
Experimental verification of theoretical light sheet formation. All images in Figure 2 show light sheet from side. (2A) Theoretical interference pattern at cylindrical lens focus. Image has been false-colored by ‘Fire’ lookup table (LUT) in Fiji (scale at top). Transverse (across sheet width) intensity line scan is overlaid on 2A in green. Full width of the central peak at half maximum intensity (FWHM) is predicted to measure 4.3 μm. (2B) Low-magnification image (upper) of light sheet focusing into fluorescent media. High magnification image (lower, red box) of cylindrical lens focal region. Green line indicates location of transverse line scan of measured intensity. (2C) Subset of image in lower portion of 2B (yellow box). Line scan (green line), scale, and coloration are consistent with predicted pattern in 2A. Measured FWHM of central sheet is 4.3 μm, in agreement with the theoretical prediction.

### LITE operates at native, diffraction-limited spatial resolution

The main goal of LITE microscopy is to combine the use of high-NA objectives to maximize resolution and detection efficiency with LSFM. We thus tested if LITE could be used with high-NA, oil-immersion objectives and provide the high resolution expected from those objectives. The spatial resolution of LITE images should depend solely on the objective NA and the wavelength of emitted fluorescent light. Therefore, spatial resolution in LITE images should be identical to spatial resolution in epi-illumination images, when the objective and samples are the same. In order to quantitatively test whether the spatial resolution is the same, we suspended sub-diffraction (100 nm diameter) fluorescent beads in 2% agarose and acquired images from the same field of beads using LITE (Fig. 3A) and epi-illumination (Fig. 3B) with a high-NA detection objective (60X 1.49 NA oil immersion). We then measured the point spread function (PSF) of each bead in three dimensions by fitting a Gaussian trace to pixel intensity and interpolating the FWHM in each dimension (Fig. 3C). The Gaussian FWHMs of beads visualized with LITE (blue) are identical to those visualized with epi-illumination (orange) (Fig. 3D-F). Spherical aberration artifacts in the z-resolution of the objective were identical for LITE and epi-illumination (Fig. 3F), supporting the conclusion that LITE operates at the expected resolution for the chosen objective.

**Figure 3.**
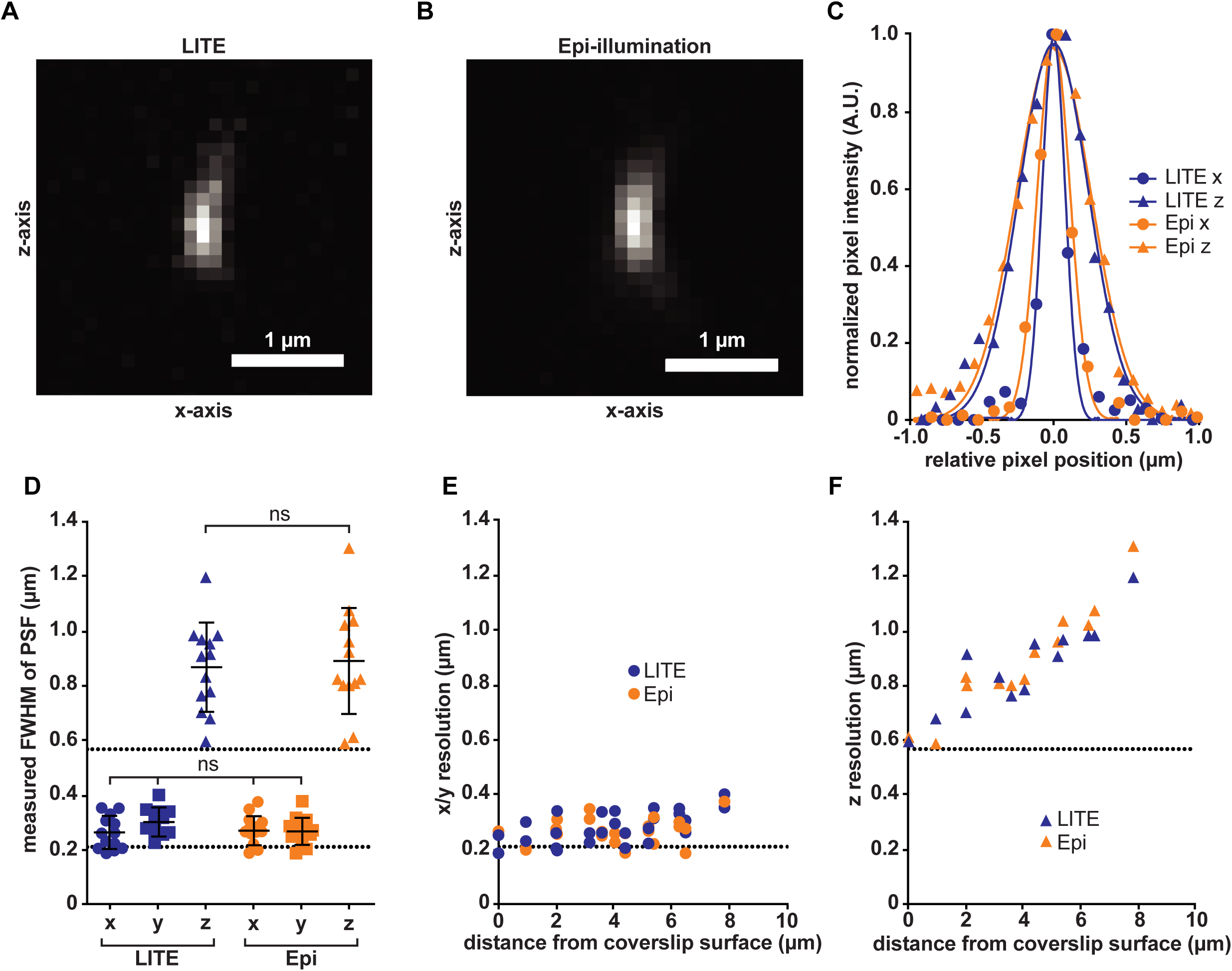
Quantification of LITE spatial resolution. (3A) Image of a fluorescent 100-nm bead, visualized using LITE. Image is maximum intensity projected along y-axis to show lateral (x) and axial (z) resolution. (3B) Image of same bead from 3A, visualized with epi-illumination. (3C) Pixel intensity values for line scans across x-axis (circles) and z-axis (triangles) for LITE (blue) and epi-illumination (orange) images of bead in 3A and 3B, respectively. Gaussian fits of intensity for each dimension are overlaid in corresponding colored lines. (3D) Plots of FWHM for Gaussian fits to fluorescence intensity of all beads (n = 12) in x (circles), y (squares), and z (triangles) dimensions for LITE (blue) and epi-illumination (orange). Statistical significance assessed by student’s t-test (ns: p > 0.1). Upper and lower black dotted lines indicate theoretical axial and lateral (0.211 and 0.568 μm, respectively) resolution for this objective. (3E) Scatterplot of each bead’s measured lateral (x/y) resolution as a function of measured distance from coverslip surface. Dotted line corresponds to predicted resolution from 3D. (3F) Scatterplot of each bead’s measured axial (z) resolution as a function of measured distance from coverslip surface. Dotted line corresponds to predicted resolution from 3D.

### LITE significantly reduces photobleaching compared to epi-illumination

As with other modalities of LSFM, the selective plane illumination of LITE microscopy is expected to reduce the photodamage experienced by live fluorescent samples. In order to quantify the photobleaching rate of LITE-illuminated samples, we imaged early (1-4 cell) *C. elegans* embryos expressing fluorescently-tagged (GFP) histone H2B (LP148 strain)^11^. To measure the true rate of GFP photobleaching without any confounding biological variables, such as new protein translation, proteolysis, and active transport of the fluorescent signal in the z-dimension, we needed a method to inhibit these biological processes. Accordingly, we immobilized the embryos by dissection into M9 nematode media + 2 mM NaN3. This treatment inhibits adenosine triphosphate (ATP) synthesis, thereby indirectly inhibiting ATP-dependent processes such as protein translation, cytoskeleton motor protein activity, and proteolysis. Thus, any decrease in the measured fluorescent signal should be due to excitation-induced photobleaching.

Fluorescent embryos were imaged under identical growth and mounting conditions using either epi-illumination or laser-illumination via LITE. The intensities of the epi-illumination field and the LITE laser were specified to generate images with similar initial starting characteristics: namely, signal-to-background ratio (SBR, qualitatively referred to as contrast) and raw integrated fluorescence density. We found that LITE preserves SBR over the course of imaging (Fig. 4A,B). Epi-illumination (orange) starts at a lower SBR and approaches the lower limit of 1.0 (a level precluding analysis) more quickly than LITE (blue; Fig. 4B).

**Figure 4.**
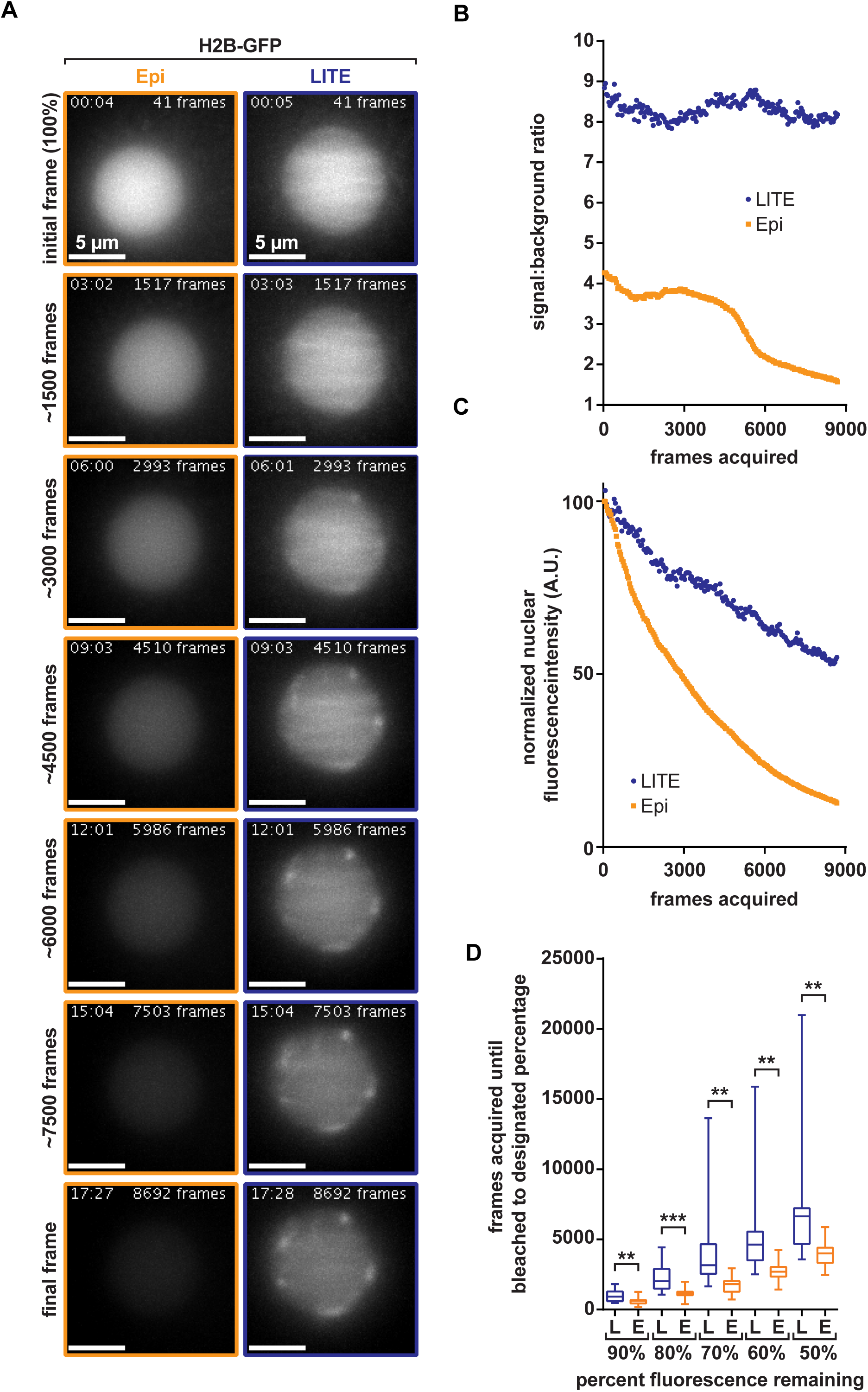
Quantification of LITE photobleaching rates. (4A) Representative image sets of *C. elegans* embryos expressing GFP-tagged histone H2B construct to visualize nuclei. Representative images show P1 nucleus. All images were taken using the same 60X 1.4 NA oil-immersion objective with a frame exposure time of 100 ms, a z-step size of 0.5 μm, a z-range of 20 μm, and no delay between timepoints. Images shown are z maximum intensity projections. Epi and LITE images are outlined in orange and blue, respectively. Lengths of time the nuclei were exposed to the laser (LITE) or arc lamp (epi) are denoted Fadero *et al*., 2017 31 in the upper left hand corner of each image in the format of minutes:seconds. Cumulative number of frames acquired up until displayed images were acquired are denoted in the upper right hand corner of each image. Rows in 4A represent images of the nuclei (internally scaled to initial frame of each nucleus) taken after denoted number of frames (left of rows). (4B) Measured signal-to-background ratios of LITE (blue) and epi (orange) nuclei shown in 4A. (4C) The raw integrated density values of the nuclear regions-of-interest (ROIs) for LITE (blue circles) and epi (orange squares) image sets represented in (4A). (4D) Box-and-whiskers plots of all nuclei (n ≥ 16), illustrating the number of frames acquired before nuclei bleached to 90, 80, 70, 60, and 50% intensities for both LITE (blue boxes, ‘**L**’) and epi (orange boxes, ‘**E**’). LITE significantly increases the number of frames that can be acquired before fluorescent nuclei bleach to 90, 80, 70, 60, and 50% of their original intensities (p < 0.01).

In addition to preserving SBR, LITE also decreases the rate at which the fluorescent signal photobleaches. At equivalent frame numbers, the nucleus visualized with LITE is brighter than that visualized with epi-illumination (Fig. 4C). We quantified the fluorescence intensities of nuclei over time and found that the detected fluorescence decreased more rapidly in the epi nucleus (orange) than in the LITE nucleus (blue; Fig. 4C). In order to more thoroughly illustrate the photobleaching improvement from epi-illumination to LITE, we measured the number of frames we could acquire from nuclei before the samples bleached to 90, 80, 70, 60, or 50% (Fig. 4D, S5). On average, LITE significantly increases the number of frames that can be acquired before the nuclei have bleached to a given percent starting intensity. In sum, compared with epi-illumination, LITE preserves SBR and reduces photobleaching.

### LITE is compatible with a variety of fluorescent organisms

These data suggest that LITE microscopy imparts less photodamage onto fluorescent samples while maintaining the resolution and detection efficiency to which cell biologists are traditionally accustomed, a novel combination of benefits that has not yet been achieved by other LSFM modalities. To demonstrate the utility of LITE microscopy with any coverslip-mounted biological sample, we imaged six popular model organisms with various fluorescent markers (Fig. 5A-E). We selected one plant, *Arabidopsis thaliana* (Fig. 5C; Movie S6), three animals, *Caenorhabditis elegans* (Fig. 5A; Movie S7), *Drosophila melanogaster* (Fig. 5D; Movie S8), and *Hypsibius dujardini* (Fig. 5E; Movie S9), one mammalian cell culture line, HeLa cells (Fig. 5B; Movie S10), and one fungus, *Ashbya gossypii* (Fig. 6; Movie S11) to illustrate the broad phylogenetic spectrum of modern model organisms accessible by LITE. These organisms also exhibit a wide range of sizes, from ∼30 μm (HeLa, Fig. 5B) to ∼1 cm (*A. thaliana* seedlings, Fig. 5C) in maximal length. In *C. elegans* expressing a fluorescently-tagged kinesin (LP447), chromosomes could be resolved between centrosomes (Fig. 5A; Movie S10). In a human cultured cell expressing a fluorescently-tagged kinetochore protein, no phototoxic effects (such as cell cycle arrest) were observed over 28,826 frames (122 minutes; Fig. 5B and inset). In plant cells, SAUR63-YFP decorates the cell membrane and fine intracellular structures, which we can visualize in three dimensions with little-to-no out-of-focus autofluorescence (Fig. 5C; Movie S6). In *Drosophila*, punctate and junction-associated fluorescently tagged Axin was observed during embryonic germband extension, which occurred at a normal rate with no detectible photobleaching despite exposure for 22,724 consecutive frames (130 minutes; Fig. 5D and inset, Movie S8). LITE is also compatible with imaging of fixed, fluorescently stained samples such as an adult tardigrade, where staining of actin (green) and the outer cuticle (magenta) reveal the intricate network of muscle fibers (Fig. 5E; Movie S9). Collectively, these data reveal that LITE can be used to visualize these organisms at high native resolution with constant illumination (i.e. no laser shuttering) for over two hours without any observable phototoxic effects (Fig. 5B,D).

**Figure 5.**
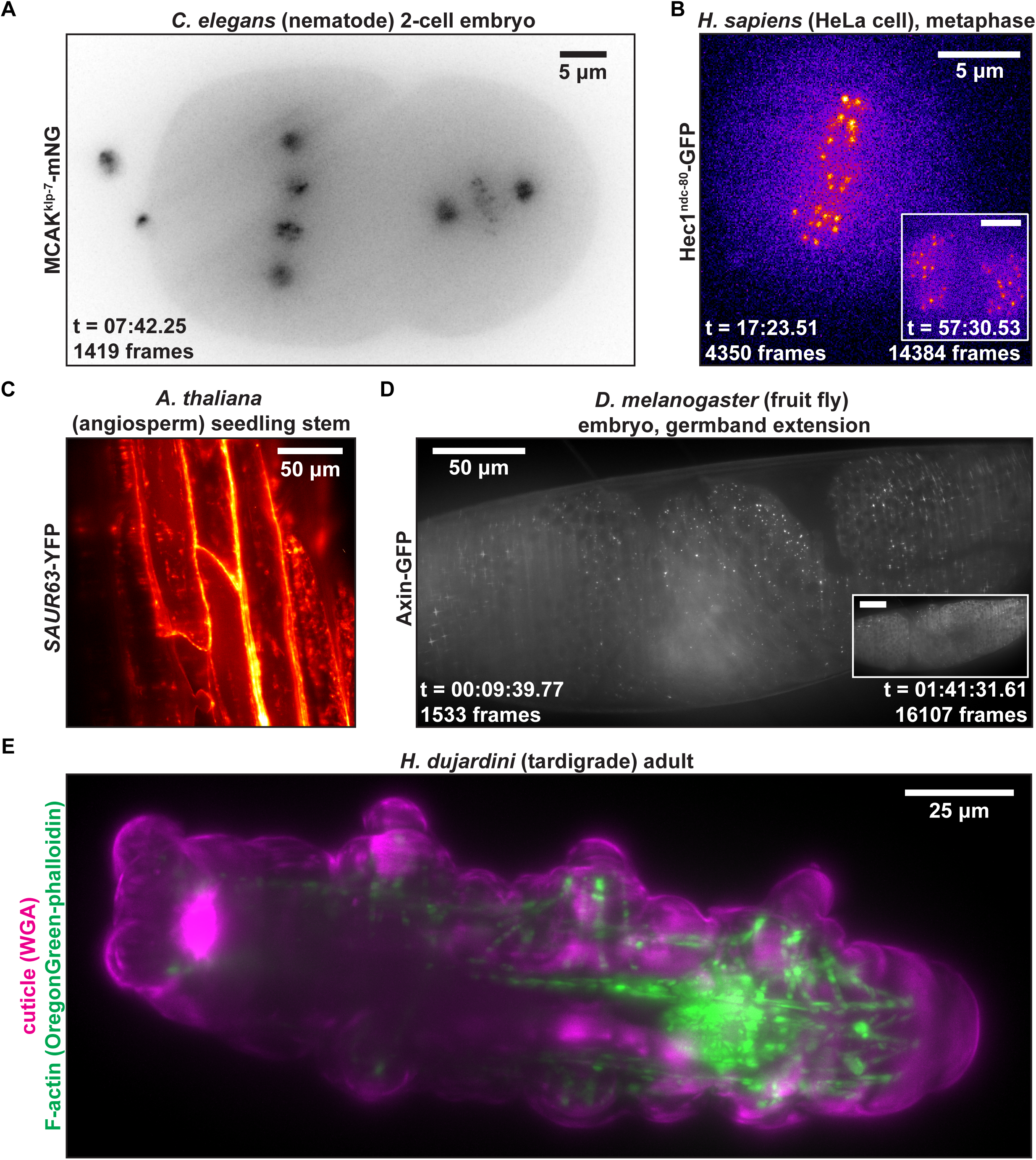
Representative LITE fluorescent images taken of a variety of model organisms, including *C. elegans* (5A), *H. sapiens* (5B), *A. thaliana* (5C), *D. melanogaster* (5D), and *H. dujardini* (5E). Fluorescent constructs imaged in each organism are delineated to left of each representative image. Images presented in 5A, 5B, and 5D are taken from the full movies available in Movies S7, S8, and S10, respectively. Images in 5C and 5E are static images taken from three-dimensional z-stacks, which are presented fully in Movies S9 and S11, respectively. Insets in 5B and 5D show images taken from later timepoints (identically scaled) to show low photobleaching. All two-dimensional images presented in Fig. 5 are maximum intensity projections of a z-series.

**Figure 6.**
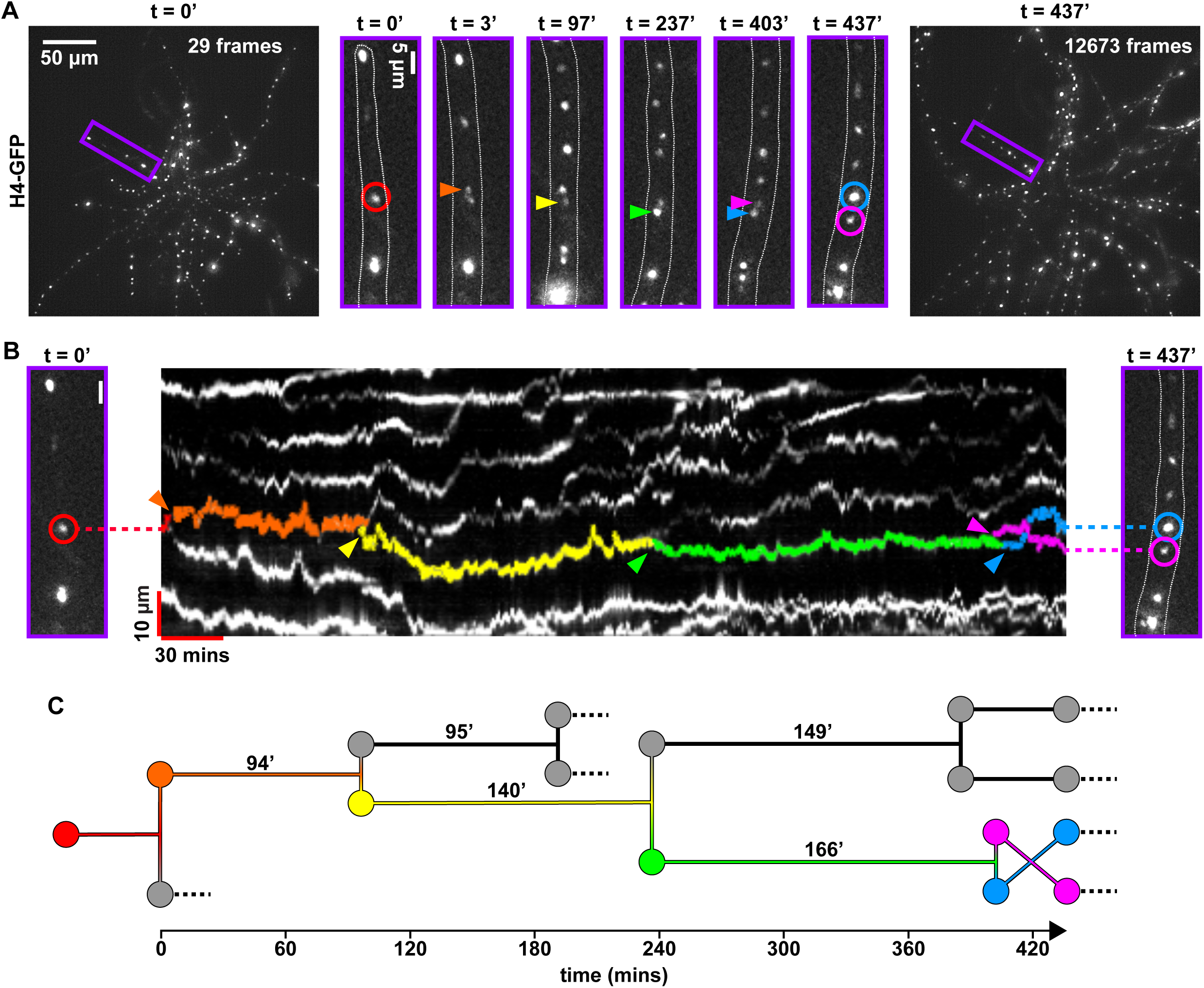
Long-term nuclear pedigrees in *A. gossypii*. (6A) Initial (left image), final (right image), and selected subsets (purple boxes, middle six images) of a 7-hour timelapse of *A. gossypii* nuclei. Purple box in left-hand image (t = 0’) is magnified and displayed to the right, outlined in purple. Outlines of hypha are shown as dotted white lines in each subset. Red circle denotes parent (1^st^ generation) nucleus to be tracked throughout timelapse. Images shown at 3’, 97’, 237’, and 403’ denote times at which mitotic division events (anaphase or telophase) of parent nucleus (3’) or its descendants (97’, 237’, 403’, 437’). Birth events (mitoses) of tracked nuclei in images are denoted with orange, yellow, green, and blue/pink arrows, respectively. Image at 437’ denotes final image of hypha acquired during the timelapse, with blue and pink arrows denoting location of fifth-generation nuclei. All images in Figure 6 were taken with a 60X 1.4 NA oil-immersion objective with a 500 ms exposure per frame, 0.5 μm z-steps, and a 14 μm z-range. Images were deconvolved with 8 iterations of a Richardson-Lucy deconvolution algorithm. (6B) Kymograph of region-of-interest (ROI) around hypha from 6A. Multicolored arrows denote same events as in 6A. Tracks of nuclei have been false-colored to highlight their lifespans, with birth of each colored nucleus denoted with colored arrowheads. Note the nuclear bypassing event of blue and pink nuclei at ∼415’. (6C) Nuclear pedigree tree of the lineage highlighted in 6A and 6B. Colored nuclei correspond to the colored arrows shown in 6A and 6B, and colored tracks correspond to the false-colored tracks in 6B. Cell cycle lengths (as measured by the length of time between mitoses) are indicated in minutes above each nuclear lifespan. Dotted lines indicate nuclei that moved out of the ROI, could not be tracked, or underwent their next division outside of the acquisition timeline.

### Long-term imaging with LITE enables nuclear lineage analysis

We next set out to demonstrate the power of combining long-term timelapse imaging with low photodamage and high spatiotemporal resolution. The filamentous fungus *Ashbya gossypii* has emerged as a powerful system in which to study syncytial cell biology^12^. Despite existing in a common cytoplasm, *Ashbya* nuclei proceed through the cell cycle out-of-sync with each other. Previous statistical analyses investigating the source of nuclear asynchrony in *Ashbya* have been based only on single pairs of sister nuclei born of a single mitotic event^13,14^, limiting robust statistical analysis of division patterns across multiple generations. However, long nuclear cycles (between 40 and 200 minutes)^14^ and highly oscillatory nuclear motions^15^ in *Ashbya* necessitate high spatiotemporal resolution, 4D imaging for at least two iterations of the average nuclear cycle (∼three hours) to trace lineages across multiple nuclear generations. To date, tracking nuclei for this duration at high spatiotemporal resolution has been confounded by photobleaching and phototoxicity. To overcome these limitations, we used LITE to image *Ashbya* and track nuclear motion and mitotic asynchrony continuously for over seven hours. We expressed a fluorescent histone (H4-EGFP) in *Ashbya* to detect nuclei for measuring motion and division (Fig. 6A; Movie S11). Nuclear divisions in a hypha (Fig. 6A, purple box) were readily identified and tracked for five generations (colored arrows). After 437 minutes of imaging, the *Ashbya* cell was alive and not detectably photobleached (Fig. 6A,B; Movie S11). Kymograph analysis enables us to create a temporally scaled pedigree of the nuclear generations (Fig. 6B,C). In sum, LITE is a powerful approach for long-term, high spatiotemporal resolution live imaging.

## Discussion

Traditionally, LSFM has been used to reduce photodamage to fluorescent samples by reducing the illumination to only the focal volume of the detection objective, but its geometry has limited use of high-NA objective lenses. LITE is the first SPIM/LSFM modality that allows the use of any objective, allowing researchers to take full advantage of the efficiency of high-NA objectives. If, for example, LITE is used with a 1.49 NA oil-immersion objective, this setup accepts 88% more emitted fluorescence and offers a 26% increase in native lateral resolution (Fig. 3) than the 1.1 NA water-dipping objective currently used with the Lattice Light Sheet^5^. Collecting more light affords LITE the ability to generate brighter images, which in turn allows the user to illuminate the sample with proportionally less intense laser power to collect the same number of photons as with other SPIM/LSFM modalities, which in turn lowers the photobleaching rate (Fig. 4). The high native spatial resolution of LITE (Fig. 3) will allow cell biologists to obtain images with the spatial resolution to which they are accustomed without sacrificing (and, likely improving) temporal resolution, since LITE does not require deconvolution of multiple structured views as does structured illumination microscopy (SIM).

LITE is also compatible with several other common aspects of modern microscopy. LITE can be installed non-obtrusively on any upright or inverted stand, allowing for the use of standard equipment, such as eyepieces, objective turrets, and trans-illumination (Fig. S3). In addition, since the native point-spread function of LITE is identical to that of epi-illumination (Fig. 3), standard post-acquisition deconvolution algorithms can be used on LITE images just as with epi-illumination. For example, the Richardson-Lucy deconvolution algorithm was used for our *Ashbya* images to increase the contrast between the nuclei and the cytoplasm.

LITE is less photodamaging than epi-illumination, both in the rate at which fluorophores photobleach (Fig. 4C) and the preservation of the image contrast over the acquisition time (Fig. 4B). By selectively illuminating a thin slice of the sample (Fig. 2), LITE reduces the background (a combination of out-of-focus signal and out-of-focus autofluorescence) relative to the in-focus signal, thus increasing the overall image SBR. High SBR provides high contrast of the structure of interest from the confounding out-of-focus background fluorescence, as well as from sample autofluorescence. The higher variability in the LITE photobleaching rates (Fig. 4D) could be attributed to variability in sheet alignment, chamber construction (Fig. S2-A), or biological noise. Although the LITE sheet measured 4.3 μm thick FWHM (Fig. 2D), in actual cellular imaging conditions, it behaved as if it were thinner. Due to the complexities of cells, this phenomenon is difficult to measure and is best illustrated by the observation that focusing the detection objective (without moving the sheet) by ∼1 μm resulted in being outside the excitation volume. While we have no experimental evidence for this observation, it is conceivable that, given the sheet has a Gaussian intensity profile, only the very peak of the focal volume contains a photon density adequate for fluorophore excitation. Regardless of the variability in sheet alignment or its functional thickness in living samples, our work demonstrates that LITE can be used to image fluorescent samples for longer periods of time than with epi-illumination (Fig. 4D; Movie S4).

As has been observed with current LSFM designs^3-6^, we found that LITE decreases the fluorophore bleaching rate in comparison to epi-illumination (Fig. 4). Theoretically, this decrease could allow users to reach an equilibrium between photobleaching and turnover at a higher signal and higher SBR with LITE than with epi-illumination. Furthermore, we observed an intriguing phenomenon in several of our model organisms in which fluorescence intensity does not detectably decrease over the course of the timelapse (Fig. 5B, 5D, 6; Movies S8, S10, S11). To explain this phenomenon, we suggest that addition of new fluorophores in live organisms could compensate for loss via photobleaching. If the translation, maturation, and loading of unbleached biological fluorophores collectively result in a simple linear increase in fluorescence, fluorophore turnover could compensate for most photobleaching in live-cell fluorescence microscopy, provided the photobleaching rate is low enough. Understanding this phenomenon will require further study, as it requires characterization of protein abundance and turnover rates to accurately calculate the photobleaching rate in living, developing samples.

We are confident that the decreased rate of photobleaching that LITE offers will allow cell biologists to observe intracellular dynamics at higher native spatiotemporal resolution and for significantly longer periods of time than previously possible using other modes of fluorescence microscopy. We have demonstrated one application of LITE in tracing nuclear lineages (Fig. 6). Lineage tracing has powerful implications, as asymmetric and symmetric inheritance of factors that determine cellular behavior is integral in determining how cells born of a single ancestor can differentiate to different fates.

In the past, we have used the model fungal system *Ashbya gossypii* where divide asynchronously in a common cytoplasm^13^. Previous work has found that individual nuclear cycles in a single *Ashbya* cell can vary significantly in their timing^13^, suggesting that there exists nuclear-intrinsic and/or -extrinsic factors that influence nuclear timing. A limitation of past nuclear tracking experiments^14,15^ was that photobleaching and phototoxicity prevented long-term imaging that would allow collection of nuclear lineage data over multiple generations, limiting the ability to robustly test for lineage-dependent similarities in nuclear timing.

With LITE microscopy, we are now able to image nuclei for over seven hours to visualize multiple rounds of nuclear division with no noticeable photodamaging effects (Fig. 6; Movie S11). These data will allow us to study the heritability of division timing over several generations and further our understanding of how heritable nuclear-intrinsic signals contribute to division asynchrony in *Ashbya*. These sorts of extended image series and statistical analyses are relevant to establishing lineages and division patterns in any cell type, from stem cells to tissues.

Beyond tracking nuclei in *Ashbya*, we demonstrate that LITE can be effectively used to visualize fluorescent labels in a wide variety of organisms at high native spatial resolution. With LITE, cell biologists can now image without photodamage far longer than with conventional modes of fluorescence microscopy. In addition, cell biologists can reduce the photodamage to their samples without sacrificing spatial resolution or detection efficiency. Thus, LITE allows biologists to observe practically any live, fluorescent organism with unprecedented efficiency and resolution for previously unattainable periods of time. These newfound observations using LITE will undoubtedly allow biologists to better understand the intricacies of cellular and subcellular dynamics.

## Acknowledgements

The authors would like to thank Dr. Edward Salmon for helpful advice, equipment, and HeLa cell lines. We also thank Dr. Ilya Golub for advice on photomask design and placement, Dr. Jason Reed and Dr. Punita Nagpal for *Arabidopsis* samples and generating Movie S9, all members of the Maddox laboratories for helpful discussions and advice, Dr. Dan Dickinson for helpful discussion and advice, and the UNC-Chapel Hill Kenan Makerspace for 3D printing. This work was funded by the NSF CAREER Award #1652512, NSF Award #1616661, NIH R01 #5R01GM102390-03, and the 2016 UNC-CH Biology Departmental Grant.

## Author contributions

T.C.F. and P.S.M. co-designed and assembled all LITE prototypes, including the final prototype (Fig. S1, S3). T.C.F. and P.S.M. co-formulated general LITE equations presented in Methods sections 1-4, designed Chamber A (Fig. S2A), and co-performed the experiments represented in Fig. 1, 2, 3. T.C.F. and P.S.M. co-drafted the Abstract, Introduction, Methods (1-4, 6, 8), and Results sections and Figures (1-5, S1, S3, S4) of the manuscript. K.R. maintained *C. elegans* strains, performed the experiments in Figures 4, S4, and Movie S4, and co-drafted Figures 4, S4, and Movie S4 with T.C.F. and P.S.M. T.M.G. and A.S.G. co-maintained *A. gossypii* stocks. T.M.G. and T.C.F. co-imaged *A. gossypii* in Figure 6 and Movie S11, co-drafted the discussion section, and co-drafted Figure 6 and Movie S11. A.S. maintained HeLa cell lines and co-drafted Methods section 7. A.S. and T.C.F co-imaged HeLa cells in Figure 5B and Movie S10. M.D., T.M.G., N.L.A., A.S.G., A.S.M., P.S.M., and T.C.F. co-designed Chamber B. M.D., N.L.A., and T.C.F. co-drafted Methods section 7, and co-drafted Figure S2. K.N.S. and M.P. co-maintained *D. melanogaster* lines. K.N.S. and T.C.F. co-imaged *D. melanogaster* for Figures 5D and Movie S8, co-drafted Methods section 7, and co-drafted Figure 5D and Movie S8. J.K.H. and B.G. co-maintained *C. elegans* strains for Figure 5A and Movie S7. J.K.H. and T.C.F. co-imaged *C. elegans* for Figure 5A and Movie S7, co-drafted Methods section 7, and co-drafted Figure 5A and Movie S7. T.C.B. maintained, fixed, and stained *H. dujardini* cultures. T.C.B. and T.C.F. co-imaged *H. dujardini* specimens for Figure 5E and Movie S9 and co-drafted Methods section 7 and Figure 5E and Movie S9. A.S.M. and P.S.M. provided all funding for design and construction of the LITE prototypes.

## References

1. Stelzer, E.H.K., Cremer, C., and Lindek, S. Theory and Applications of Confocal Theta Microscopy. Zoo. Stud. 34 (1), 67–69 (1995).

2. Laissue, P.P. et al. Assessing phototoxicity in live fluorescence imaging. Nature Methods, 657–661 (2017).

3. Huisken, J. et al. Optical sectioning deep inside live embryos by selective plane illumination microscopy. Science 305 (5686), 1007–1009 (2004).

4. Santi, PA. Light sheet fluorescence microscopy: A review. J. Histochem. Cytochem. 59 (2), 129–138 (2011).

5. Chen, B.C. et al. Lattice light-sheet microscopy: imaging molecules to embryos at high spatiotemporal Simultaneous Multiview capture and fusion improves spatial resolution in wide-field and light-sheet microscopy. Optica 3 (8), 897–910 (2016).

6. Wu, Y. et al. Simultaneous Multiview capture and fusion improves spatial resolution in wide-field and light-sheet microscopy. Optica 3(8), 897–910(2016).

7. Golub, I., Chebbi, B., & Golub, J. Toward the optical “magic carpet”: reducing the divergence of a light sheet below the diffraction limit. Opt. Lett. 40 (21), 5121–5124 (2015).

8. Pai, JH et al. Photoresist with Low Fluorescence for Bioanalytical Applications. Anal. Chem. 79 (22), 8774–8780 (2007).

9. Murashige, T., and Skoog, F. A revised medium for rapid growth and bioassays with tobacco tissue cultures. Physiol. Plant., 473–497 (1962).

10. Smith, F.W. and Jockusch, E.L. The metameric pattern of Hypsibius dujardini(Eutardigrada) and its relationship to that of other panarthropods. Front. in Zoo. 11 (66),2014.

11. Dickinson et al. Engineering the Caenorhabditis elegans genome using Cas9-triggered homologous recombination. Nat. Meth. 10 (10), 1028–1034 (2013).

12. Roberts, S.E., and Gladfelter, A.S. Nuclear autonomy in multinucleate fungi. Curr. Opin. Microbiol., 60–65 (2015).

13. Gladfelter, A.S., Hungerbuehler, K., and Philippsen, P. Asynchronous nuclear division cycles in multinucleated cells. Journ. Cell. Biol. 172 (3): 347–362 (2006).

14. Nair, D.R. et al. A conserved G1 regulatory circuit promotes asynchronous Fadero et al., 2017 36 behavior of nuclei sharing a common cytoplasm. Cell Cyc. 9 (18), 3771–3779 (2010).

15. Anderson, A.C. et al. Nuclear Repulsion Enables Division Autonomy in a Single Cytoplasm. Curr. Bio. 23 (20), 1999–2010 (2013).

